# Neural mechanisms of sequential dependence in time perception: The impact of prior task and memory processing

**DOI:** 10.1101/2023.05.07.538104

**Authors:** Si Cheng (程思), Siyi Chen (陈思佚), Stefan Glasauer, Daniel Keeser, Zhuanghua Shi (施壮华)

## Abstract

Our perception and decision-making are susceptible to prior context. Such sequential dependence has been extensively studied in the visual domain, but less is known about its impact on time perception. Moreover, there are ongoing debates about whether these sequential biases occur at the perceptual stage or during subsequent post-perceptual processing. Using functional Magnetic Resonance Imaging (fMRI), we investigated neural mechanisms underlying temporal sequential dependence and the role of action in time judgments across trials. Participants performed a timing task where they had to remember the duration of green coherent motion and were cued to either actively reproduce its duration or simply view it passively. We found that sequential biases in time perception were only evident when the preceding task involved active duration reproduction. Merely encoding a prior duration without reproduction failed to induce such biases. Neurally, we observed activation in networks associated with timing, such as striato-thalamo-cortical circuits, and performance monitoring networks, particularly when a “Response” trial was anticipated. Importantly, the hippocampus showed sensitivity to these sequential biases, and its activation negatively correlated with the individual’s sequential bias following active reproduction trials. These findings highlight the significant role of memory networks in shaping time-related sequential biases at the post-perceptual stages.

**Significance Statement:** Our study explores the neural mechanisms of sequential dependence in time perception and reveals that active reproduction of time duration in the previous trial can bias subsequent estimates, resulting in a sequential dependence effect. In contrast, passive viewing of a stimulus without reproducing its duration does not produce this effect. At the neural level, we observed increased activity in memory regions like the hippocampus when sequential biases were reduced. Furthermore, we found a negative correlation between hippocampal activation and sequential bias following active reproduction trials, suggesting that the involvement of memory networks mediates how we are influenced by past experiences when judging time.

## Introduction

The world around us is relatively stable and predictable over a short period. A traffic sign at a crossroad will turn periodically into red and green in a predictable way. During rush hour, a traffic jam is likely followed by another. Our experience is thus useful to guide our decisions because the past and the present often correlate. Research has demonstrated that our current perception is biased by recent events, referred to as serial dependence or sequential dependence (Holland and Lockhead 1968; Cross 1973; Fischer and Whitney 2014; Cicchini et al. 2018; Glasauer and Shi 2022; Pascucci et al. 2023). Such sequential bias has primarily been explored in the context of non-temporal features, such as orientation, color, and motion direction, particularly with a recent paradigm highlighting the influences of the difference between the current and previous stimuli on perceptual biases (e.g., Fischer and Whitney 2014; Cicchini et al. 2017; Bae and Luck 2020; Barbosa and Compte 2020).

There is ongoing debate regarding the underlying mechanisms of sequential dependence. Two main perspectives have emerged. The first view proposes that sequential dependence is thought to maintain perceptual stability and continuity by integrating past and current information to filter out abrupt noises, serving as a perceptual mechanism (Liberman et al. 2016; Whitney et al. 2022). The second view connects sequential dependence to prior task and response-related post-perceptual factors (Fritsche et al. 2017; Pascucci et al. 2019; Bae and Luck 2020; Ceylan et al. 2021). For instance, sequential dependence is only observed when the current and previous tasks are the orientation judgments (Bae and Luck 2020), indicating that encoding the previous stimulus was not sufficient but the task-related response in previous trials was essential for sequential effects to occur.

In this latter view, working memory plays a crucial role in sequential dependence, as it involves integrating preceding stimuli with current sensory inputs for decision-making and motor plans (Bliss et al. 2017; Kiyonaga, Scimeca, et al. 2017; Bae and Luck 2019; Fornaciai and Park 2020a). Studies have shown the sequential effect increases when the memory retention interval increases (Bliss et al. 2017), and decreases when the short-term memory maintenance in the premotor cortex is interrupted with TMS (de Azevedo Neto and Bartels 2021). A recent fMRI study showed that neural activity in low-level V1 is opposite to behavioral sequential dependence, which suggests that the effect emerges in high-level memory or decision-making circuits (Sheehan and Serences 2022). Moreover, a recent electroencephalogram (EEG) study used the auditory pitch categorization task and decoded neural representations of multiple features (i.e., pitch, category choice, motor response) of the current trial as well as the neural response from past-trial features on the current trial, and it found that past-trial features kept their respective identities in memory and were only reactivated by the corresponding features in the current trial, giving rise to sequential biases (Zhang and Luo 2023).

While behavioral studies reliably demonstrate sequential dependence, much of the existing literature has primarily focused on non-temporal spatial features, such as orientation, that have a circular distribution (e.g., Fischer and Whitney 2014). One advantage of using this is that it allows for a clear separation of sequential effects from pervasive central tendency and range effects (Vierordt 1868; Teghtsoonian and Teghtsoonian 1978; Shi, Church, et al. 2013; Petzschner et al. 2015; Glasauer and Shi 2021) – phenomena that are commonplace in magnitude estimations, such as duration judgments. For example, perceived durations can skew toward a mean duration derived from recent history or sampled durations (Nakajima et al. 1992; Burr et al. 2009; Jazayeri and Shadlen 2010), leading to underestimate long durations and overestimate short ones. However, only a handful of recent behavioral studies have explored trial-to-trial sequential effects on timing (Wiener et al. 2014; Wehrman et al. 2020; Togoli et al. 2021; Glasauer and Shi 2022), and even fewer have linked these effects to specific electroencephalogram (EEG) signatures (Damsma et al. 2021; Fornaciai et al. 2023). This leaves a significant gap in our understanding of neural mechanisms at play.

In this study, we employed a classic post-cueing paradigm used in sequential dependence research (Czoschke et al. 2019; Bae and Luck 2020) to investigate the neural mechanisms underlying sequential dependence in a duration reproduction task (Shi, Ganzenmüller, et al. 2013; Shi et al. 2022; Zang et al. 2022) in conjunction with MRI scanning. The task consisted of an encoding phase and a subsequent phase that was either for reproduction or passive-viewing, contingent on a post-cue that indicated “Response” or “No-response”. During the encoding phase, participants had to remember the stimulus duration, and then either reproduce it or passively observe it, as instructed by the cue. With “Response” and “No-response” trials randomly intermixed, participants had to recall the durations accurately in each case. This design allowed us to compare how passive viewing and active reproduction during preceding trials influenced the processing of the subsequent stimuli, thereby shedding light on the post-perceptual factors contributing to sequential dependence.

To preview our findings, we found behaviorally that the central tendency bias operated independently of the preceding task. However, the sequential dependence was significant only when the preceding task required active reproduction rather than passive viewing. At the neural level, our data link striato-thalamo-cortical and performance monitoring networks to time perception and prior tasks, respectively. Notably, we observed that hippocampus activity was directly linked with the sequential bias on both prior tasks and prior duration. This hippocampal activation was particularly evident during the encoding phase following passive viewing trials and led to a decrease in sequential bias. These findings highlight the involvement of post-perceptual stages that link sensory representations to responses and underscore the critical role of active timing-related and memory networks in the temporal sequential dependence.

## Materials and Methods

### Participants

21 participants (9 females; 12 males; age: Mean ± SD = 27.24 ± 3.83, range: 23–33 years) were recruited for the two-session MRI experiment^1^. All of them were right-handed and had normal or corrected-to-normal vision and color vision, no history of neurological, psychiatric, or medical disorders, and no symptoms of COVID-19 in the past two weeks. The sample size was determined based on previous studies (Fischer and Whitney 2014; Bae and Luck 2020) that usually found large effect sizes (Cohen’s *d* > 0.75), and with a power of 80% (1 − β), which yielded a minimum of 19 participants according to G*Power (Erdfelder et al. 1996). Participants provided their informed consent prior to the experiment and were compensated for their participation for 15 Euro/hour. The study was approved by the ethics committees of the Psychology Department at LMU Munich.

### Experimental design and procedure

We adopted a duration reproduction paradigm (Shi, Ganzenmüller, et al. 2013; Shi et al. 2022; Zang et al. 2022), consisting of an encoding phase and a reproduction phase (see Figure 1). Participants laid down comfortably with their head in the head coil, inside some cushions to fixate the head position and avoid motion. Participants viewed the back projector canvas (diagonal 30 inches) via an adjustable mirror positioned on top of the head coil, with a viewing distance of 110 cm. The task was presented by an MRI-compatible ProPIXX DLP LED projector (Pixx Technologies Inc, Canada).

**Figure 1.**
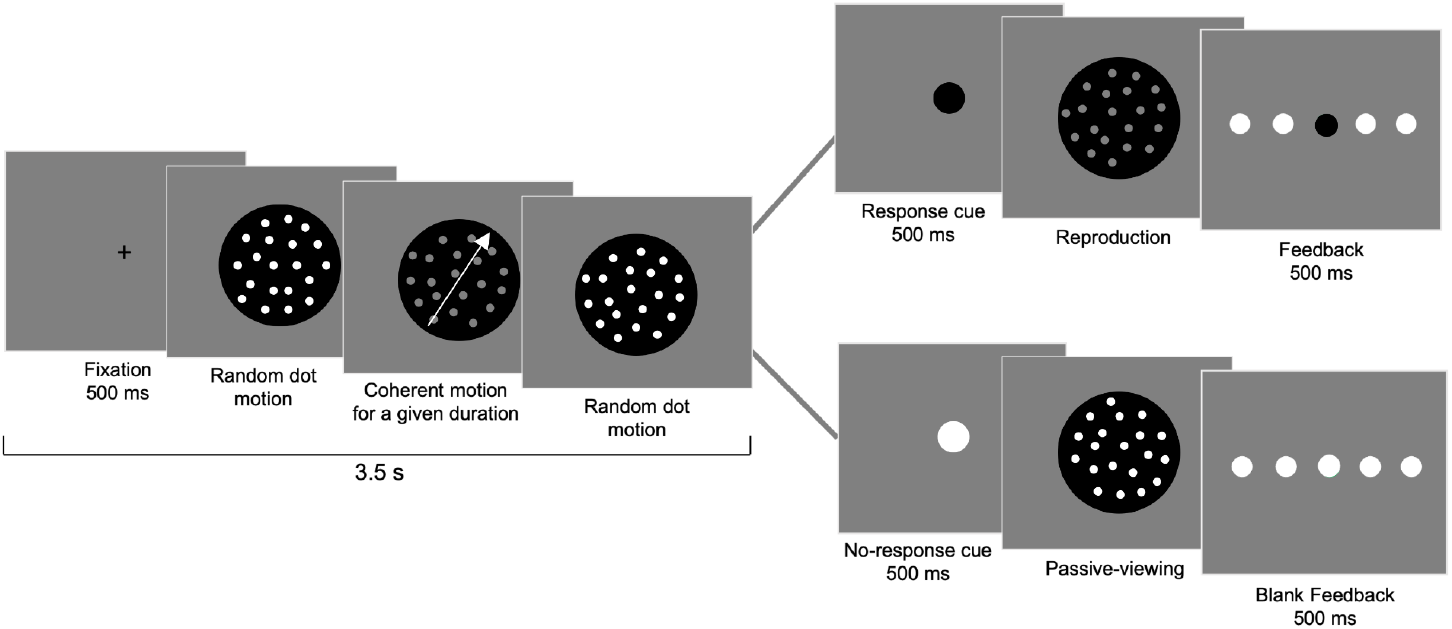
Experimental procedure. Each trial consisted of two phases: the duration encoding phase and the duration reproduction or passive-viewing phase, with the latter being contingent upon the Response / No-response cue. The trial started with a fixation, followed by the display of white random dot motion, which then changed to green dots moving coherently in one direction (e.g., the gray dots in the illustration represented the green coherently moving dots in the direction of the white arrow, not shown in the actual experiment) for a given duration before reverting back to white random dot motion. The entire presentation of the dots display lasted for 3000 ms. Participants were instructed to remember the duration of the green coherent motion. A cue, either a green (“Response” trial, represented as the black disk in the illustration) or a white (“No-response” trial) disk, was then shown on the screen, indicating the beginning of the second phase. In the “Response” trials, the green random dot motion appeared (represented as the gray dots in the illustration), and participants had to press a button when they thought the elapsed duration was the same as the perceived duration in the encoding phase. During “No-response” trials, participants only passively viewed the white random dot motion for the same duration as the coherent movement of the green dots in the encoding phase and were not required to respond. In the end, there was a feedback display featuring five horizontal white disks, with one disk changing color to indicate the accuracy of the reproduction for the “Response” trials.

Each trial began with a black cross (visual angle of 0.5°) at the center of the display on a light gray background for 500 ms, prompting participants to maintain their fixation on the center. Following this, a tunnel view display (subtended 17.8° visual angle) of white randomly moving dots (20 dots with each diameter of 0.4°, moving at a speed of 1°/s) appeared for a randomly selected duration between 400 to 600 ms. The white dots then changed to green and moved 100% coherently with a speed of 6°/s in one direction selected between 0 to 360° for a given duration, chosen from 0.8, 1.0, 1.2, 1.4, and 1.6 secs. Immediately after the given duration, the color of the dots changed back to white and moved randomly again at a speed of 1°/s. When a dot exited the tunnel view, it was regenerated at a random location within the view to maintain dot density. The entire presentation of the tunnel view lasted for 3000 ms. Participants were instructed to remember the duration of the green coherent motion.

Following the encoding phase, a cue was presented in the center for 500 ms that indicates whether participants should perform a reproduction task (“Response”) or not (“No-response”). The cue was a color disk (subtended 1.2° visual angle), which could be either green (indicating a Response trial: reproducing the duration) or white (indicating a No-response trial: doing nothing, only passive viewing). For Response trials, immediately following the offset of the cue display, a green random dot motion in a tunnel view (20 green dots with each of 0.4° at a speed of 1°/s) appeared in the center of the screen. Participants had to monitor the elapsed time and press an MRI-compatible ResponsePixx button (Pixx Technologies Inc, Canada) with their right thumb when they perceived the elapsed time as being the same as the duration of coherent movement in the encoding phase. Following the response, a feedback display appeared indicating the response accuracy with a color dot in a horizontal dot array. Each highlighted dot from the left to right five dots corresponded to a range of relative reproduction errors (error/duration): The left most represented below –30%, while the second to fifth dot indicated errors between [-30%, –5%], (–5%, 5%), [5%, 30%], or greater than 30%, respectively. The middle dot was highlighted in green to indicate an accurate reproduction, while the 2nd and 4th dots were highlighted in light red, the utmost 1st and 5th dots in dark red, with the color intensity reflecting the degree of error. The feedback lasted for 500 ms. For the No-response trial, the cue display was followed by a white random dot motion in a tunnel view (20 white dots with each diameter of 0.4°, moving at a speed of 1°/s), lasting the same amount of time as the coherent movement of the green dots in the encoding phase, and participants didn’t need to respond and just passively watched the display. Afterwards, a blank feedback display appeared for 500 ms to equate the time with the feedback display in the Response trial. The next trial started after a random interval of 2000 to 3000 ms.

Since we were interested in the impact of the preceding task on the present reproduction, and to ensure equal transitional probability from a “No-response to Response” trial and from a “Response to Response” trial while minimizing the number of trials required for MRI scanning (within one hour), we excluded the possibility of a “No-response to No-response” trial. This yielded a 2:1 ratio of the Response vs. No-response trials while maintaining equal transitional probability (see a similar approach Czoschke et al. 2019).

Prior to the formal scanning, participants received a practice block of 30 trials to familiarize themselves with the task in a sound-reduced and moderately lit chamber, near the scanning room. In the formal MRI study, each participant completed 12 blocks, with each block of 30 trials, among them 20 Response trials and 10 No-response trials. After each block, there was a short 6-second break, and after 6 blocks (comprising a session) a long 7-minute break. During this long break, participants took a rest inside the scanner, and a T1 image was acquired. The whole experiment lasted approximately 60 minutes.

### MRI data acquisition and preprocessing

All MRI data were acquired using a 3-Tesla Siemens Prisma MRI scanner (Siemens, Erlangen, Germany), equipped with a 32-channel head coil. Functional MRI images were obtained using a blood oxygenation level-dependent (BOLD) contrast-sensitive gradient-echo EPI sequence. A total of 3000 to 3300 volumes of fMRI data, depending on the duration of the experiment, were acquired for each participant through two sessions. The following parameters were used: TR = 1000 ms, multi-band factor = 4, TE = 30 ms, flip angle = 45°, FOV = 210 × 210 mm, voxels size = 3 × 3 × 3 mm, slices number = 48, slice thickness = 3 mm. In addition, structural MRI images (T1 weighted) were acquired from the sagittal plane using a three dimensional magnetization prepared rapid gradient-echo (MPRAGE) pulse sequence with the following scanning parameters: TR = 2500 ms, TE = 2.22 ms, flip angle = 8°, FOV = 256 × 256 mm, voxel size = 0.8 × 0.8 × 0.8 mm, slice thickness = 0.8 mm.

The analyses and visualization of imaging data were performed with SPM12 (Ashburner et al. 2014) and Nilearn – 0.9.2 (Abraham et al. 2014). The functional images were first preprocessed with realignment, reslicing, and slice time correction. Then, the head movement correction was done by using affine transformation in a two-pass procedure and aligning individual functional images to their mean image using 2nd-degree B-spline interpolation. Participants who had head movements more than 3 mm or rotations greater than 3° were excluded from further analysis, which yielded the exclusion of six participants in total, five for large head motion and one for T1 image distortion. We then replaced them with six new participants. The mean image of each participant was then spatially normalized to a 3 mm standard Montreal Neurological Institute (MNI) template using the “unified segmentation” approach, and the resulting deformation field was applied to the individual functional images. The normalized fMRI images were then smoothed with a 6 mm full-width-at-half-maximum (FWHM) Gaussian kernel to improve the signal-to-noise ratio and compensate for residual anatomical variations.

### Statistical analyses

#### Behavioral analysis

The reproduction errors, the difference between the reproduced duration and the physical duration, were calculated for the Response trials. To exclude trials with large reproduction errors due to accidental button presses or inattention, we applied the three-sigma rule to exclude those outliers. Then, we categorized the remaining Response trials into two types depending on the preceding task:

a. Response-Response (RR) trials that were preceded by a reproduction task, and (b) NoResponse-Response (NR) trials that were preceded by a passive viewing task.

Given that time intervals are an open scale and their judgments are subjective to both central tendency and sequential biases, we estimated the central tendency and sequential dependence effects separately. The central tendency effect results from the integration of the prior (*D_p_*) and the current duration (*D_n_*), and can be estimated through linear regression when the prior is considered fixed(Cicchini et al. 2012; Shi, Church, et al. 2013):

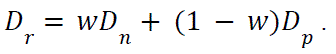

Here we applied a linear regression to find the slope (*w*), and designated 1 − *w* as the index for central tendency. An index of 0 indicates no central tendency, while an index of 1 signals a strong central tendency.

The conventional measures of serial dependence effect, which compare the current response error to the difference between the previous and the current stimuli (e.g., Fischer and Whitney 2014; Bliss et al. 2017; Kiyonaga, Manassi, et al. 2017; Cicchini et al. 2018), are not sufficient to separate sequential dependence from the central tendency bias (for more details, see Glasauer and Shi 2022). Instead, we adopted a classical approach to sequential effects (Holland and Lockhead 1968), which correlates the current error with the previous duration. However, this method could still capture a general bias, such as systematic over– or underestimation, in addition to the sequential trend. Thus, in the analysis, we only focused on the linear trends (i.e., the slopes), rather than the intercepts.

Specifically, we applied linear regression to relate the current response errors (*E_n_*) with the previous sample duration (*D_n-_*_1_), and took the slope *b* of this linear fit as the index of the sequential bias:

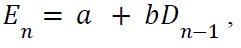

A positive slope indicates that the current error is assimilated towards the previous duration, while the negative slope suggests a repulsion from the previous one. Additionally, as a sanity check and for further verification, we also computed the sequential effect for the durations presented in future trials (n+1).

For statistical analyses, we applied linear regression, simple *t*-tests and linear mixed models according to the data structure.

#### fMRI statistical analysis

There were three types of inter-trial transitions: No-response to Response (NR), Response to Response (RR), and Response to No-response (RN). Given that the third type (RN) trials yielded no behavioral response, our analysis focused on the Response trials (i.e., RR and NR). To boost the power of the first-level fMRI analyses, we grouped the preceding durations into two categories: the “Short” and the “Long” categories. The “Short” category included durations of 0.8 and 1.0 s, and the “Long” categories were 1.4 and 1.6 s, with the middle duration of 1.2 s being excluded. Therefore, a combination of the factors of the Prior Task (RR or NR) and the Prior Duration (Short or Long) resulted in four conditions.

At the first-level analysis with individual participants, BOLD responses obtained from the smoothed images were time-locked to the onset of the target duration and modeled using a canonical hemodynamic response function (HRF) with a box-car function to represent the duration of the coherent movement. Our analysis defined four conditions through the combination of the Prior Task and the Prior Duration, each represented by a separate main canonical hemodynamic response function (HRF) regressor. To identify brain regions associated with sequential biases, we incorporated an additional parametric regressor for each main regressor. Parametric modulation serves as an index of the relationship between neural activity and the normalized response error, helping to pinpoint regions where activity varies based on specific variables (Penny et al. 2011; Mumford 2015).

Given that the current response error inherited the central tendency biases and the systematic general biases, we can not directly use them as parametric regressors for identifying neural activities associated with the sequential effect. Instead, we employed normalized errors:

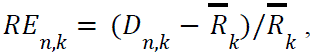

where 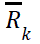 is the mean reproduction of the duration *D_k_*, *D*_n,*k*_ the reproduced duration at trial *n* of a given Duration *D_k_*, R*E*_n,*k*_ the normalized relative error. The normalized errors retained the trend of sequential dependence associated with the previous duration, yet they were free from the general bias and the central tendency linked to the current duration (see Appendix Figures S1 and S2). This approach allowed us to use it as an additional parametric regressor on the neural activity of the current trial to isolate the impact of sequential dependence. Together, we had four main regressors and four parametric regressors. The beta value of the parametric regressor reveals the strength and linear relationship between brain activity and the relative error. A positive parametric value indicates that brain activity correlated positively with the relative error, while a negative value suggests an inverse correlation between brain activity and the relative error. Importantly, as BOLD activations were analyzed during the encoding phase and sequential biases were assessed during the late reproduction phase, the parametric modulation serves to quantify both the extent and direction of how fluctuations in brain activity influence behavioral sequential bias. Additionally, six head movement parameters were added as nuisance regressors to control head motion artifacts (Lund et al. 2005). The data were high-pass filtered at 1/128 Hz. For each subject, 8 condition-specific contrast images were created (for each trial type and parametric modulator).

The respective contrast images for the main HRF regressors and the parametric regressors were subjected to the second-level analysis with flexible factorial design, separately. In the flexible factorial design, Prior Task and Prior Duration were created as within-subject factors and Participant as a random factor. The 2 (Prior Task: Response (RR) vs. No-response (NR)) × 2 (Prior Duration: Short vs. Long) ANOVAs allowed us to determine the unique contributions of each factor to the brain activity, and how they might interact with each other to affect neural processing. We conducted a whole-brain analysis to determine the candidate brain regions involved in the main effects of Prior Task and Prior Duration, as well as their interaction, by using planned t-contrasts. All contrasts were thresholded at *p* < .001, with FWE cluster correction at *p* < .05. We were interested in brain regions that showed sensitivity to the modulation of brain activity by the normalized relative deviation. Parametric estimates were extracted from the statistically significant clusters and averaged across the voxels in the individual-level analysis. Once the regions of interest (ROIs) were identified, we proceeded with a comprehensive analysis where all durations were taken into account.

To gain further insight into activation patterns, a separate general linear model (GLM) was applied to the fMRI time series, similar to the model designed above, except all individual durations were included (0.8, 1.0, 1.2, 1.4, and 1.6 s). Thus, we have 2 (Prior Tasks) × 5 (Prior Durations), 10 conditions in total. Each condition had one main HRF regressor and one parametric regressor using the normalized relative error. Beta values of individual main regressor as well as parametric modulator were calculated from the significant voxels detected above (a sphere with a diameter of 10 mm) for individual participants. To assess differences in beta values, we employed a linear mixed model analysis, incorporating Prior Task and Prior Duration as the fixed effect, and Subject as the random factor. Linear mixed models are resilient to violations of sphericity and help mitigate the risk of Type I errors (Singmann and Kellen 2019). The *p*-values reported for the mixed models were calculated using the maximum likelihood estimation.

We then conducted Spearman’s correlation (α = 0.05, two-tailed) to assess the relationship between the magnitude of the sequential dependence effect in behavior and the BOLD activity of interest. The slope of the linear regression with the normalized relative error depending on the previous sample duration was operationalized to indicate the magnitude of the behavioral sequential dependence effect. The BOLD response was measured as the r values of the main regressors in the designated regions detected above.

## Results

### Behavioral results

The reproduction errors were calculated for the “Response” trials. Overall, reproduced durations were close to the probe durations, resulting in a mean reproduction error of 35 ms. We excluded trials with large reproduction errors and the outliers were generally rare, on average only 0.89%, ranging individually from 0 to 5 outlier trials. The remaining “Response” trials were categorized into two types depending on the preceding trial: (a) “Response” to “Response” (RR) trials that were preceded by a reproduction trial, and (b) “No-response” to “Response” (NR) trials that were preceded by a passive viewing trial. Given that duration reproduction is affected by both the central tendency effect and the sequential dependence (Glasauer and Shi 2022), we estimated the central tendency and serial dependence effects separately.

#### The central tendency effect

We applied a linear regression of the duration reproduction (*D_r_*) on the current durations (*D_n_*) to obtain the slope (*w*), and used 1 − *w* as the central tendency index, with 0 indicating no central tendency and 1 strong central tendency (Cicchini et al. 2012; Shi, Church, et al. 2013). The results, shown in Figure 2a, indicate a strong central tendency effect – participants overestimated short durations and underestimated long durations. The central tendency effect was quantified through linear regression, revealing significant central tendency biases with mean central tendency index (1 − *w*) of 0.540 (*t*_(20)_ = 9.13, *p* <.001, *d* = 1.99) and 0.594 (*t*_(20)_ = 9.14, *p* <.001, *d* = 1.99) for the NR and RR respectively. But the central tendency biases were comparable between the two conditions (*t*_(20)_ = 1.748, *p* = .096, *BF_10_* = 0.830). Additionally, there was a minor positive general bias (M ± SE: 35 ± 14 ms, *t*_(20)_ = 2.452, *p* = .024, *BF_10_* = 2.494), which was comparable between the two conditions (*t*_(20)_ = 1.298, *p* = .209, *BF_10_* = 0.475). A linear mixed model with Prior Task and Current Duration as the fixed effects and Subject as the random factor also confirmed similar results: no effect of Prior Task (Coef = –0.012, 95% *CI* [-0.038, 0.014], *p* = .380), but a significant main effect of Current Duration (Coef = 0.406, 95% *CI* [0.341, 0.471], *p* < .001). And their interaction was not significant (Coef = 0.054, 95% *CI* [-0.038, 0.146], *p* = .251). That is, the central tendency and the general bias were not influenced by the prior trial task. The lack of a significant difference in the central tendency effect between preceding task types might be primarily due to the same distribution and range of the tested durations for two prior tasks, yielding a consistent long-term representation of the prior durations across tasks. This agrees with previous findings that randomly mixing durations leads to a generalization of the prior across conditions (Roach et al. 2017).

**Figure 2.**
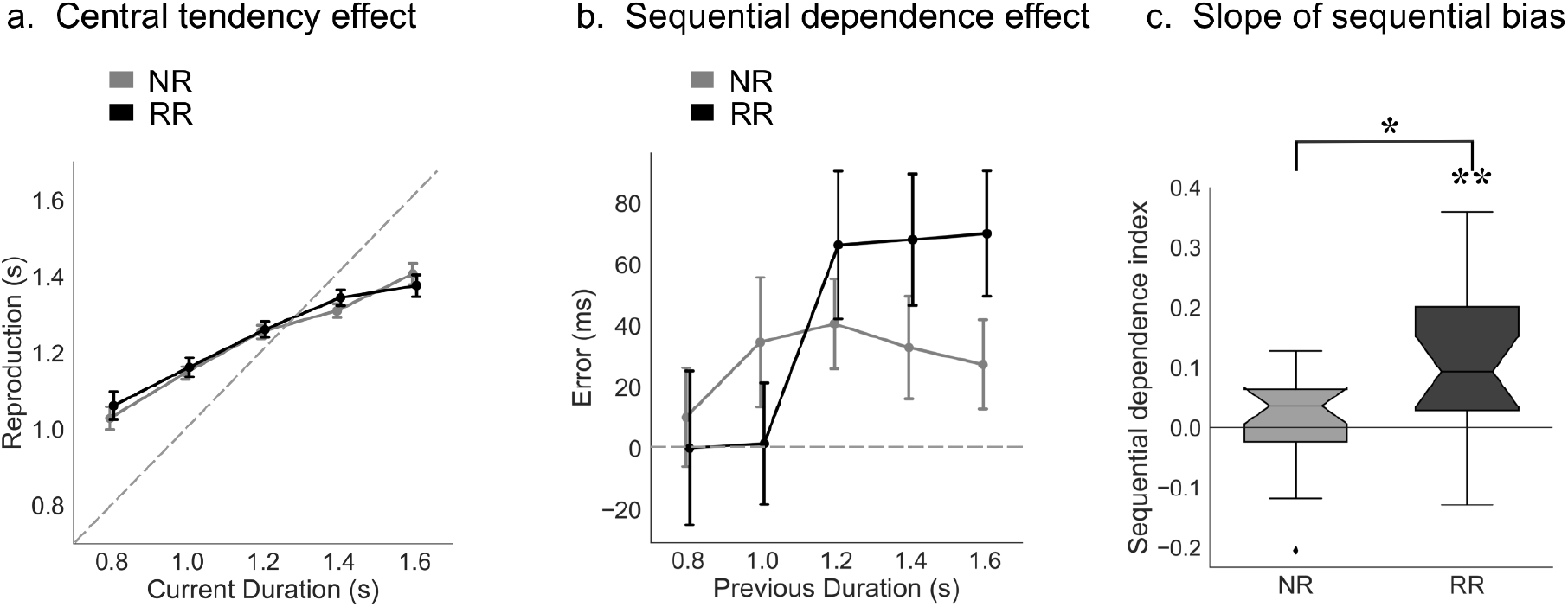
(**a**) The central tendency effect. The duration reproductions are shown as a function of the actual duration, separated for the inter-trial type: No-response/Response (NR) and Response/Response (RR) trials. Error bars represent ± SEM. (**b**) The sequential dependence effect. The response errors are plotted on the durations from previous trials, separated for NR and RR conditions. Error bars represent ± SEM. (**c**) Notched boxplots of the sequential dependence slope for NR and RR conditions. The box plot depicts the sequential dependence effect, measured by the slope, for each condition. The top and bottom of the notched box represent the interquartile range (between 25% and 75%), and the notch in the box is the 95% confidence interval for the median; whiskers exclude outliers. \**p* <.05, \*\**p* <.01.

#### The sequential dependence effect

Figure 2b shows that the current response errors increased with increasing prior durations, suggesting a sequential dependence effect. This was captured by the slopes of the linear regression, which was only significantly positive for the RR condition (*b* = 0.103, *t*_(20)_ = 3.668, *p* = .002, *BF_10_* = 25.043), but not for the NR condition (*b* = 0.017, *t*_(20)_ = 0.922, *p* = .368, *BF_10_* = 0.332). Additionally, the slope for the RR condition was significantly larger than that in the NR condition (*t*_(20)_ = 3.056, *p* = .006, *BF_10_* = 7.510, see Figure 2c). The results were further confirmed by the linear mixed model with Prior Task and Prior Duration as the fixed effects and Subject as the random factor. Neither the main effect of the Prior Task (Coef = 0.012, 95% *CI* [-0.005, 0.030], *p* = .174) nor the Prior Duration (Coef = 0.016, 95% *CI* [-0.027, 0.060], *p* = .464) were significant. However, their interaction effect (Coef = 0.087, 95% *CI* [0.025, 0.149], *p* = .006) was significant, indicating a notable difference in the slopes between the NR and RR conditions. These results suggest that active reproduction in the preceding trial increased the sequential dependence on the current reproduction, whereas passive viewing did not. To avoid statistical artifacts (Cicchini et al. 2014), we also tested the sequential dependence effect on the durations presented in future trials (n+1), which showed no significance (*ps* > .216). Our behavioral results provide clear evidence that merely passively perceiving an interval is not sufficient to bias subsequent duration estimates.

### fMRI Results

#### Whole-brain analysis

The main analysis was focused here on the effects of prior tasks on the current duration reproduction (i.e., RR and NR). The combination of the factors of the Prior Task (RR or NR) and the Prior Duration (Short or Long) resulted in four conditions, each represented by a separate main HRF regressor and an accompanying parametric regressor incorporating the normalized relative error R*E_i,k_*. The results from the main HRF regressors reflect different brain activations across different conditions, while the findings of the parametric regressors reflect the covariate changes of brain activities to the normalized relative errors in duration reproduction. In the following subsections, we report the results separately for the main HRF regressors and parametric regressors.

#### Main HRF regressors

We conducted contrast analyses using 2 (RR vs. NR) × 2 (Prior Short vs. Prior Long) within-subject ANOVAs. Table 1 and Figures 3a and 3b show the significant regions identified through this contrast analysis. The results show that the right inferior frontal gyrus (RIFG), a region associated with response inhibition (Aron et al. 2007; Hampshire et al. 2010), exhibited greater activation during RR trials compared to NR trials. Conversely, the left precuneus and a large cluster comprising the left precentral gyrus, left postcentral gyrus, the right posterior-medial frontal and the right thalamus were more active during NR trials compared to RR trials.

**Figure 3.**
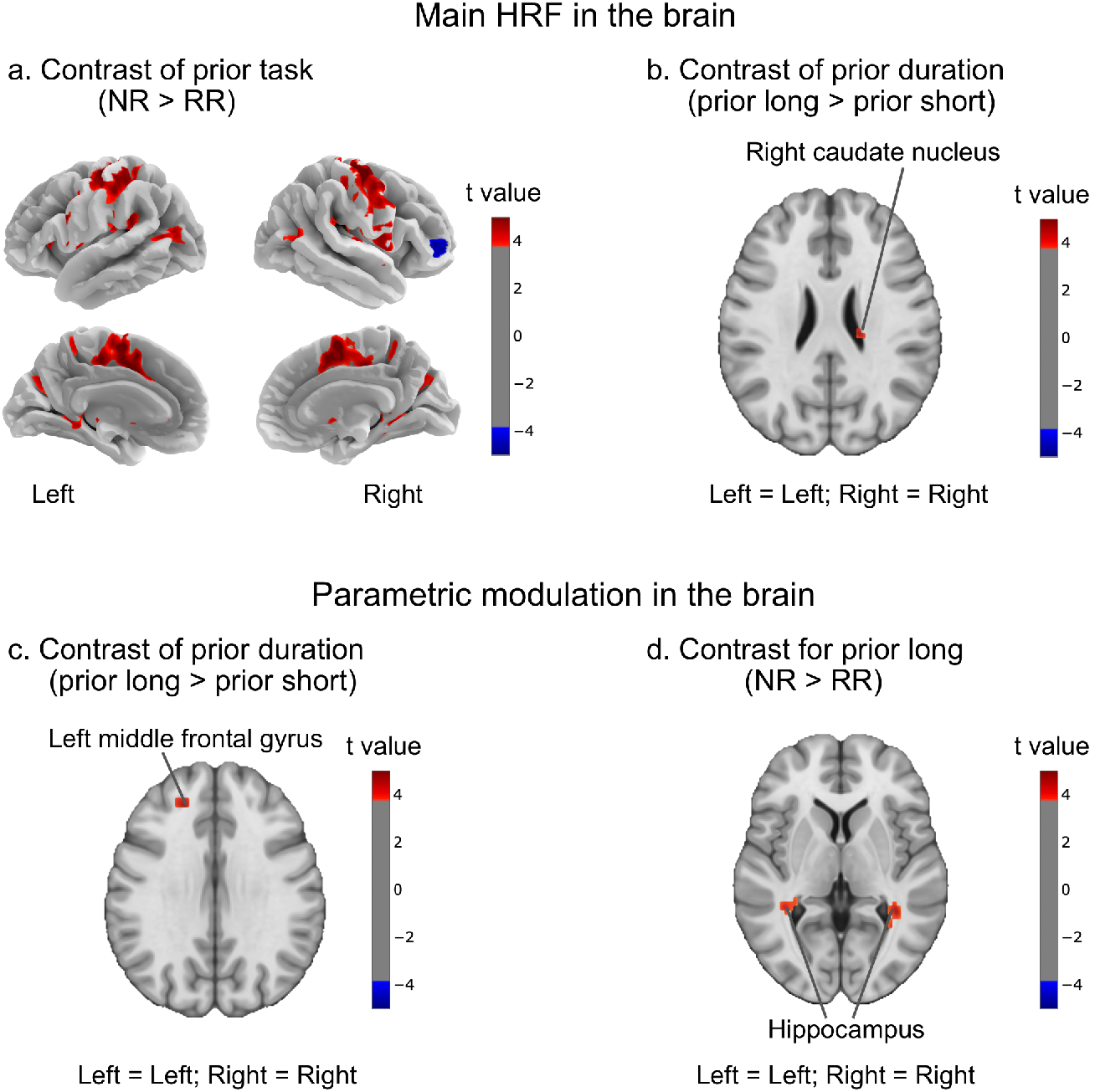
Whole-brain activation results. (**a**) and (**b**) are brain regions activated in the main HRF. (**a**) Whole-brain activation patterns colored in red-blue reflected invoked BOLD signals driven mainly by the prior task. The red-marked regions (the left precentral gyrus, the left postcentral gyrus, the left precuneus, the right posterior-medial frontal, and the right thalamus) were activated more for the preceding No-response (NR) task as compared to the preceding Response task (RR), while the blue-marked region (right inferior frontal gyrus) was activated more for the RR than the NR condition. (**b**) Whole-brain activation patterns colored in red-blue reflected invoked BOLD signals that were driven by the prior duration. The red-marked region (right caudate nucleus) was activated more for the long duration as compared to the short duration. (**c**) and (**d**) are brain regions that show sensitivity to normalized relative error in different conditions. (**c**) Main effect of prior duration for the parametric estimates with the normalized error. The red-marked region (left middle frontal gyrus, MNI coordinates: –21, 38, 32) was activated more with a larger normalized relative error for the preceding long duration than the short duration. (**d**) The normalized error-dependent differences between prior tasks (NR vs. RR) in the prior long-duration condition were expressed in the hippocampus (MNI coordinates: –36, –37, –7; MNI coordinates: 39, –31, –7). All thresholding was done at *p* < .05 with FWE-corrected at the clustered level. Neurological convention was used (Left = Left and Right = Right).

**Table 1.**
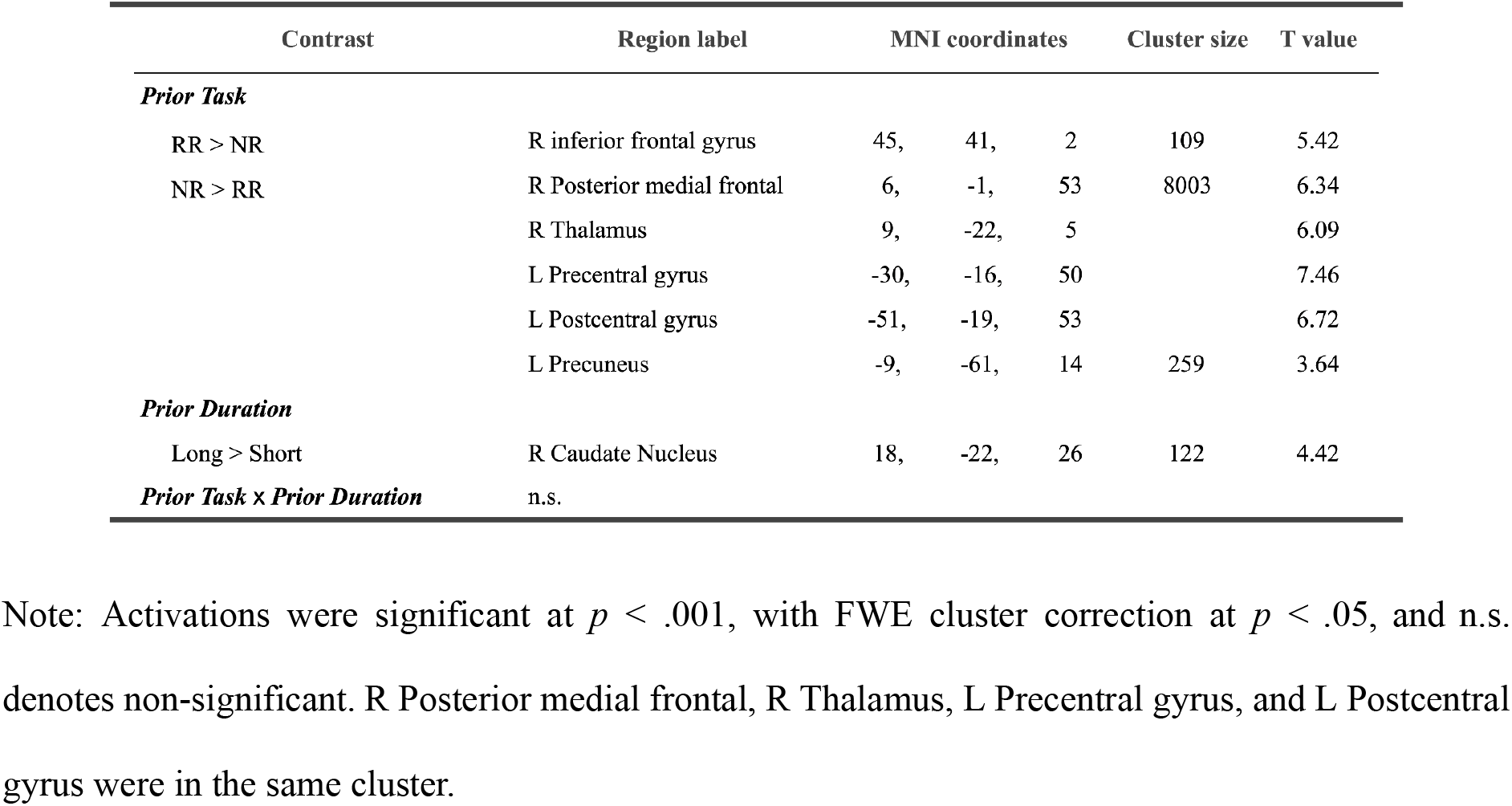
Activations associated with the main HRF regressors defined by the two factors Prior Task and Prior Duration.

The RIFG is a critical region for response inhibition and detecting important cues (Aron et al. 2007; Hampshire et al. 2010). The activation of the RIFG for the contrast RR vs. NR is likely due to a Response trial (R) can be followed by Response or No-response trials which require response inhibition. In contrast, for the contrast NR vs. RR, we observed high activation in cognitive control and performance monitoring networks, including the right posterior-medial frontal cortex (Debener et al. 2005) and the left precentral gyrus and left postcentral gyrus, as well as the left precuneus (Ridderinkhof et al. 2004; Fitzgerald et al. 2010). Increased activation in the posterior-medial frontal cortex (Ridderinkhof et al. 2004; Fitzgerald et al. 2010) is believed to be engaged in the cognitive control and performance monitoring that leads to response selection and performance adjustments in subsequent trials (Debener et al. 2005), while the precentral gyrus is the site of the primary motor cortex and the postcentral gyrus which constitutes the primary somatosensory cortex also takes part in motor control of the body (Hari et al. 1998; Dijkerman and de Haan 2007). This was likely due to the activation of the primary motor cortex for those “Response” trials while not for the “No-response” trials, and the sequential structure of the NR, which indicates that the upcoming trial is “Response” before the cue appears. The thalamus plays a key role in the cortico-thalamic-basal ganglia timing circuits (Mole et al. 2018; Yin et al. 2022).

The contrast analysis conducted between the prior short and long durations revealed that the right caudate nucleus, the main component of the dorsal striatum, was more activated when the prior trial was long than short (see Table 1 and Figure 3b). The activity in the dorsal striatum (including the caudate nucleus) is closely related to the representation of temporal information in working memory (White 2009; Merchant et al. 2013; Teki and Griffiths 2016; Yin et al. 2022). Thus, our findings suggest preceding long vs. short durations could impact the working memory trace in the following trial, consistent with previous event-related fMRI studies on working memory and time interval demonstrating that activity in the caudate nucleus increased with an increasing number of intervals in the sequence (Rao et al. 2001; Coull and Nobre 2008; Teki and Griffiths 2016).

#### Parametric regressors

To investigate the modulation of sequential effects, we employed the normalized error – which is free from the general bias and the central tendency – as a parametric regressor. The results of this analysis are presented in Table 2 and Figures 3c and 3d. These results display the contrasts of the parametric regressors based on a 2 (Prior Task: Response vs. No-response) × 2 (Prior Duration: Short vs. Long) factorial analysis. By examining these contrasts, we can identify which brain regions during the encoding stage correlate with variations in normalized errors based on preceding conditions, which could help pinpoint the brain regions sensitive to sequential bias in time perception.

**Table 2.** Activations associated with the parametric regressors defined by the two factors Prior Task and Prior Duration.

| Contrast | Region label | MNI coordinates |  |  | Cluster size | T value |
| --- | --- | --- | --- | --- | --- | --- |
| *PriorTask* | n.s. |  |  |  |  |  |
| *Prior Duration* |  |  |  |  |  |  |
| Long > Short | L Middle frontal gyrus | -21, | 38, | 32 | 124 | 4.46 |
| *Prior Task x Prior Duration* |  |  |  |  |  |  |
| Long (NR > RR) | L Hippocampus | -36 | -37, | -7 | 139 | 4.09 |
|  | R Hippocampus | 39, | -31, | -7 | 305 | 5.01 |

Notes: Activations were significant at $p < .001$ , with FWE cluster correction at $p < .05$ , and n.s. denotes non-significant.

We observed a significant main effect of Prior Duration. As seen in Table 2 and Figure 3c, the left middle frontal gyrus (LMFG; MNI coordinates: –21, 38, 32, including 124 voxels) was more positively correlated with the normalized error when the duration of the preceding trial was long vs. short. This suggests that influences of the left middle frontal gyrus activation on normalized error depend on the preceding duration. Figure 4a depicts the parametric values extracted from the left middle frontal gyrus. On average, the parametric value was positive (0.412) when the preceding duration was long, but negative (–0.691) when the preceding duration was short. Specifically, activation of the left middle frontal gyrus led to a positive trend of the relative error when the preceding duration was long, but a negative trend of the relative error when the preceding trial was short. Recall the right caudate nucleus was found to activate more strongly in response to a long preceding duration compared to a short one, and the right caudate nucleus is functionally connected to the left middle frontal gyrus (Robinson et al. 2012), which is associated with working memory encoding (Nee et al. 2013; Dandolo and Schwabe 2019). Given this association, it is possible that these two regions work together to regulate inter-trial dependence in duration judgments.

**Figure 4.**
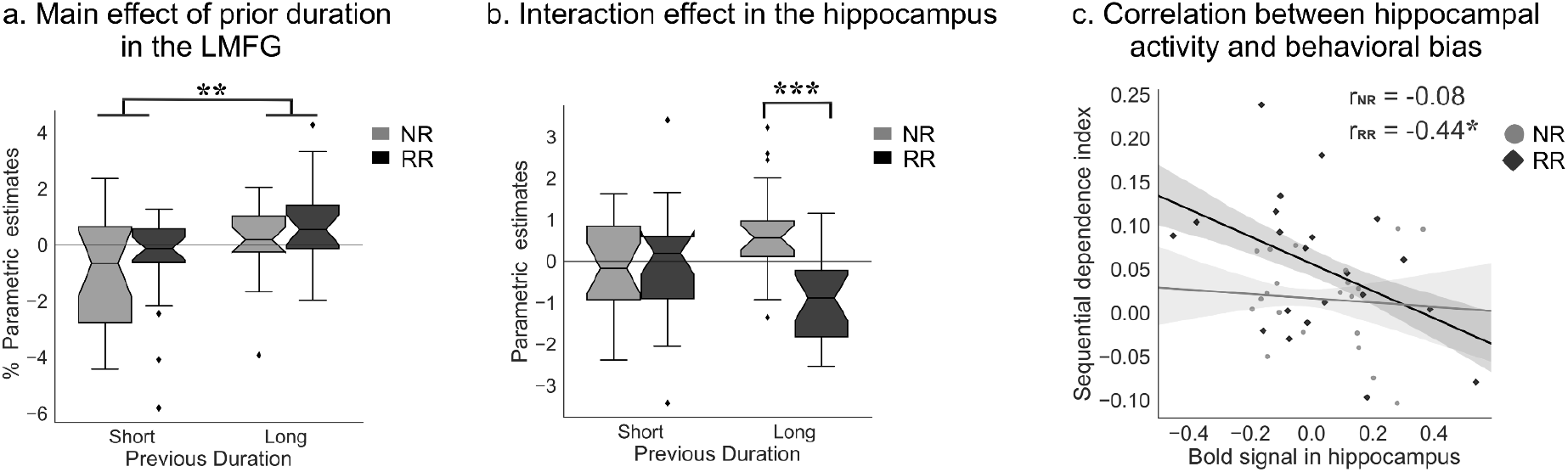
(**a**) Notched boxplots for the mean parameter estimates (r) from the left middle frontal gyrus (LMFG) for the parametric regressor for the prior tasks (NR: No-response; RR: Response), separated for the category of the previous duration (Short vs. Long). The top and bottom of the notched box represent the interquartile range (between 25% and 75%), and the notch in the box is the 95% confidence interval for the median; whiskers exclude outliers. (**b**) Notched boxplots for the mean parameter estimates (r) from the hippocampus for the parametric regressor. (**c**) Correlations between the mean BOLD signal in the bilateral hippocampus (MNI coordinates: –36, –37, –7, including 139 voxels; MNI coordinates: 39, –31, –7, including 305 voxels) and the behavioral bias (slopes of the linear regression with the normalized errors to the previous sample duration) across participants, separated for the RR and NR conditions. The correlation was significantly negative for the RR condition (r*_RR_* = –.44, *p* < .05), but not for the NR (r*_NG_* = –.08, *p* = .73). The least-square fit lines are shown. * *p* <.05, ** *p*< .01, and *** *p*< .001.

Additionally, the analysis revealed a significant Prior Duration × Prior Task interaction in the left and right hippocampus (left MNI coordinates: –36, –40, –7, including 16 voxels; right MNI coordinates: 39, –31, –7, including 25 voxels; with small-volume correction at *p* < .05 FWE corrected). Further analyses revealed the most significant contrast results from the left and right hippocampus between RR and NR conditions when the preceding trial was long, which showed different sensitivity to the normalized bias (See Figure 3d, MNI coordinates: –36, –37, –7, including 139 voxels; MNI coordinates: 39, –31, –7, including 305 voxels). As shown in Figure 4b, on average the parametric values were comparable between NR and RR when the preceding duration was short, but exhibited opposite signs for the preceding long durations. In trials preceded by a long/No-response task, the brain activity of the hippocampus was associated with a positive trend in normalized errors. Conversely, in trials preceded by a long/Response task, the hippocampal brain activity was linked to a negative trend in normalized errors.

We then further conducted a correlation analysis to examine the relationship between the hippocampal BOLD signal and the sequential dependence index. To keep the analysis consistent, we used the sequential index obtained from the normalized error. The analysis revealed a negative correlation (*r* = –.44, *p* = .044, two-tailed; Figure 4c) for RR trials (for both short and long duration), suggesting more activation in the hippocampus leads to less sequential dependence for RR trials. However, there was no correlation for NR trials (*r* = –.08, *p* = .733, two-tailed; Figure 4c), partly because the activation in the hippocampus was generally higher for NR than RR trials (Figure 5a).

**Figure 5.**
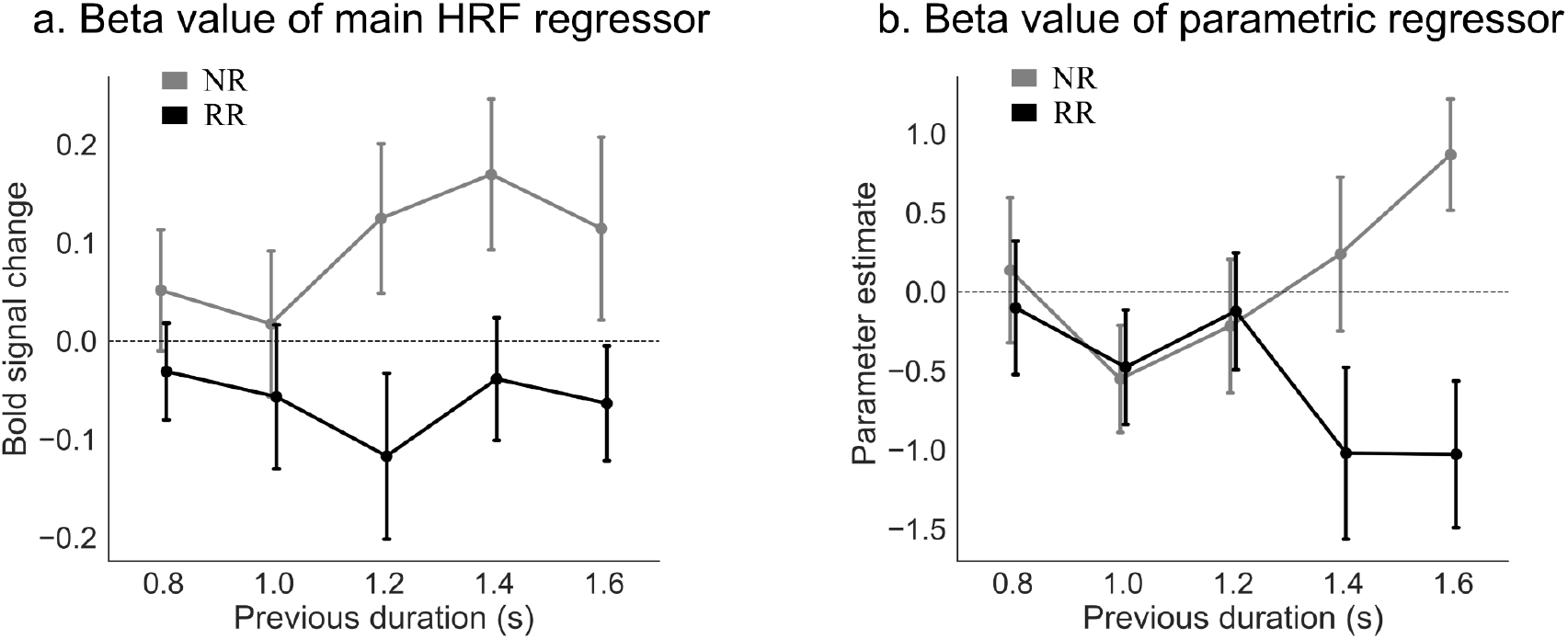
(**a**) BOLD signal change estimated from the hippocampus, plotted as a function of previous duration, separated for the previous task (NR: No-response, RR: Response). (**b**) Mean beta value of the parametric regressor extracted from the bilateral hippocampus (MNI coordinates: –36, –37, –7, including 139 voxels; MNI coordinates: 39, –31, –7, including 305 voxels) plotted as a function of previous durations, separated for the previous task (NR: No-response, RR: Response).

#### ROI analysis on the hippocampus

Note that in the whole brain analysis, we observed a large significant activated cluster (consisting of 8003 voxels) in the contrast NR > RR analysis (see Table 1), which included 0.7% (56 voxels) of this cluster that belonged to the hippocampus. To more closely examine the neural activation patterns in the bilateral hippocampus in the cluster we obtained in our ROI analysis (MNI coordinates: –36, –37, –7, including 139 voxels; MNI coordinates: 39, –31, –7, including 305 voxels), we applied a separate GLM on BOLD signals to model the previous tasks (RR vs. NR) with each of the five previous durations (0.8, 1.0, 1.2, 1.4, and 1.6 s) for the individual subject, and extracted the r values from the main and parametric regressors. The r values of the main HRF regressors are shown in Figure 5a. Further linear mixed model on the r values with the fixed factors of Prior Duration and Prior Task, and a random factor of participants revealed a significant difference between prior tasks (*b* = 0.16, 95% CI [0.08, 0.233], *p* < .001), but not among prior durations and the interaction between Prior Task and Prior Duration (all *ps* > .239). On average, trials with preceding passive viewing (i.e., NR), compared to trials with preceding active reproduction (i.e., RR), had 16% more activations in the hippocampus.

Figure 5b shows that the parametric value decreases with an increase in the previous duration for the RR condition, but increases for the NR condition. A linear mixed model on the parametric values with fixed factors of Prior Duration and Prior Task revealed a significant main effect of Prior Task (*b* = 0.64, 95% *CI* [0.12, 1.17], *p* = .016), and a significant interaction between Prior Task and Prior Duration (*b* = 2.32, 95% *CI* [0.48, 4.17], *p* = .014). The interaction was due to opposite linear trends observed between the parametric values and the previous duration for the NR and RR conditions: a positive trend for the NR (*b* = 1.13, 95% CI [-0.18, 2.43], *p* = .091), but a negative trend for the RR (*b* = –1.20, 95% CI [-2.50, 0.11], *p* = .072). Although the individual trends (the slopes of 1.13 vs. –1.20) were marginally significant from zero, the difference between the two (*b* = 2.32) was significant, particularly for the previous long durations (Figure 5b). Interestingly, for the RR condition, the negative trend of the parametric value with the previous long duration was opposite to the significant positive trend of the sequential dependence effect (Figure 2b). This implies that decreased brain activity in the hippocampus was associated with a high sequential error, possibly due to the recycling of prior information in working memory, which results in strong serial dependence (Sheehan and Serences 2022; Whitney et al. 2022).

## Discussion

This study investigated neural mechanisms that underlie serial dependence in time perception. We conducted a duration reproduction task with a post-cue to manipulate between active reproduction and passive viewing of durations, aiming to determine where the serial dependence originates. We found a strong central tendency effect in duration reproduction, regardless of the preceding task. However, the reproduction errors depended on the preceding task and duration, showing a positive serial dependence effect only for trials with consecutive reproduction (RR), but not for trials preceded by passive viewing (NR).

Our study seeks to extend upon previous research, which has primarily focused on serial dependence in non-temporal domains (Holland and Lockhead 1968; e.g., Fischer and Whitney 2014). Previous studies have shown that serial dependence requires active retrieval of a recent past (Suárez-Pinilla et al. 2018; Bae and Luck 2020; Fornaciai and Park 2020b; Ranieri et al. 2022). For instance, a study on motion direction judgment (Bae and Luck 2020) revealed that sequential dependence was only present when the preceding task was identical, as opposed to being different, such as judging the color of the motion stimuli. Our study extends this research by demonstrating that in the temporal domain, simply encoding the previous stimulus was not enough to produce a sequential effect. Employing a post-cueing paradigm offers the advantage of ensuring participants remain attentively engaged with the durations in each trial to perform the task correctly. If merely observing in the prior trials could bias subsequent duration estimates, we would expect to see some serial dependence even after just encoding from the previous trial. By contrast, our results revealed that for sequential dependence in time perception to occur, action component is essential, which suggests that the origin of this temporal sequential dependence is likely rooted in high-level, post-perceptual decisional and integrational processes (Fritsche et al. 2017; Kiyonaga, Scimeca, et al. 2017; Roach et al. 2017; Pascucci et al. 2019; Bae and Luck 2020; Ceylan et al. 2021).

It should be noted, however, that our findings do not dismiss the potential influences originating from the perceptual stage, as commonly identified in studies involving non-temporal stimuli (Fornaciai and Park 2018, 2020b; Czoschke et al. 2019; Togoli et al. 2021). Convergent evidence shows that serial dependence could emerge at different stages. For example, evidence has emerged where irrelevant stimuli, whether they are response-irrelevant inducers (Fornaciai and Park 2018) or simultaneously presented irrelevant (Czoschke et al. 2019), can either attract or repulse estimates in subsequent tasks. Intriguingly, these studies tend to position these task-irrelevant inducers close in time to the target (e.g., within a second in Togoli et al., 2021), which could promote potential perceptual integration. Well-documented phenomena, like ensemble perception (Whitney and Yamanashi Leib 2018), show a tendency to attract individual items toward the ensemble mean (Nakajima et al. 1992; Zhu et al. 2021; Baykan, Zhu, Allenmark, et al. 2023; Baykan, Zhu, Zinchenko, et al. 2023). A classic manifestation of this is the time-shrinking illusion, where successive intervals seem to blend into one another (Nakajima et al. 1992; Burr et al. 2009). From a Bayesian standpoint, the ensemble prior assimilates the target duration (Shi and Burr 2016; Zhu et al. 2021). The distinction between action and inaction can impact prior updates in unique ways. For instance, it has been shown that duration judgment exhibits a decisional carryover effect, a tendency to report the current stimulus as being similar to a prior one (Wiener et al. 2014; Wehrman et al. 2020). A study by Roach et al. (2017) highlighted that when participants were tasked with reproducing clearly delineated durations (either short or long) associated with specific spatial locations, the durations from different locations were merged together forming a single prior that significantly influenced their reproductions in all locations. However, merely passive observation of durations from one location left time reproductions from another location largely unaffected. This points to the influential role of responses, suggesting that action may serve as a common cause for temporal assimilation, leading to sequential effects predominantly in consecutive trials that demand a response. Our study using the post-cue paradigm further confirmed that the sequential effect in the time domain relies heavily on late decision stages that engage action. It is essential to highlight that in our study, since both passive no-response trials and response trials covered the same duration range, we cannot conclusively determine if central tendency would differ when passive and active trials span different ranges. Nonetheless, the response-driven common cause hypothesis does suggest a possible variation in central tendency biases. It would be interesting for further studies to validate this prediction.

At the neural level, during the duration encoding phase, BOLD signals were enhanced for NR trials relative to RR trials. This was observed within a network associated with cognitive control and response preparation (Ridderinkhof et al. 2004; Dijkerman and de Haan 2007; Fitzgerald et al. 2010), which includes regions like the right posterior-medial frontal, the left postcentral gyrus, and the left precuneus. Additionally, the right thalamus, a critical component of cortico-thalamic-basal ganglia timing circuits (Mole et al. 2018; Yin et al. 2022), also showed activation. After a passive-viewing trial with no response demands, the response preparation network and cortico-thalamic-basal ganglia timing circuits were better primed for the upcoming trial. In contrast, the right inferior frontal gyrus (RIFG), essential for response inhibition and cue detection (Aron et al. 2007; Hampshire et al. 2010; Hartwigsen et al. 2019), displayed increased activity during RR trials compared to NR trials. This enhanced activity occurs because, after an active reproduction trial, the type of next trial (Response or No-response) remains uncertain until the appearance of the post cue.

The contrast between the prior long and short durations revealed greater activation in the right caudate nucleus for the prior long duration. The caudate nucleus, a key part of the dorsal striatum, plays a critical role in the striato-thalamo-cortical network (Rao et al. 2001; Coull and Nobre 2008; Teki and Griffiths 2016). Serving as a “core timer” of the timing system (Meck et al. 2008), the caudate nucleus holds temporal “memories” in its GABAergic medium spiny neurons (MSNs) via dopamine-facilitated long-term potentiation and short-term plasticity mechanisms (Allman and Meck 2012; Kononowicz 2015). Adjustments to corticostriatal synaptic weights by MSNs in the dorsal striatum could tune them to specific time intervals encoded by coincident oscillatory patterns, increasing the likelihood of them firing upon similar intervals in the future (Kononowicz et al. 2016; Yin et al. 2022). The observed sequential dependence in the present study likely reflects these residual temporal “memories” left in the MSNs of the caudate nucleus from the previous trial.

Using parametric modulation analysis on brain activity, we unveiled compelling patterns reflecting the intricate relationship between sequential bias and BOLD activations. Specifically, the left middle frontal gyrus showed greater sensitivity to sequential bias when previously exposed to long durations as opposed to short ones, resonating with observed activations in the right caudate nucleus. It is widely recognized that the left middle frontal gyrus in concert with the fronto-striatal pathway (Darki and Klingberg 2015; Teki and Griffiths 2016) contributes to working memory encoding (Nee et al. 2013; Dandolo and Schwabe 2019). Past research has solidified this connection, showing that activity in the caudate nucleus and the frontal cortex systematically increased with an increasing number of intervals in the sequence (Rao et al. 2001; Coull and Nobre 2008; Teki and Griffiths 2016). This emphasizes the indispensable role of the fronto-striatal pathways in time perception (Matell et al. 2005). Drawing from these insights, our findings thus argue that the strong modulation of sequential bias in the left frontal gyrus can be attributed to the high involvement of working memory encoding and fronto-striatal timing circuits in processing prior long durations.

Furthermore, the hippocampus displayed a higher activity level for NR trials than for RR trials, working together with the preparation network that includes the precuneus – a region known for its crucial role in working memory (Hebscher et al. 2018; Ren et al. 2018). This high activation of the hippocampus during NR trials likely serves to actively preserve the encoded duration, effectively shielding potential sequential biases. In contrast, the subdued hippocampal activation during RR trials suggests a less active maintenance of the current duration, possibly because the information from prior trials is being reused (Sheehan and Serences 2022; Whitney et al. 2022). Additionally, the parametric modulation analysis further elucidated the hippocampus’s role in shaping sequential bias in both NR and RR conditions. Specifically, as the previous duration lengthened, the parametric value increased for the NR but decreased for the RR condition. Their difference reached significance at long durations (i.e., 1.4 and 1.6 secs). Further supporting this, we found a significant negative correlation between BOLD signals from the hippocampus and the sequential bias, but only in the RR condition. That is, reduced hippocampal activation was linked to a greater likelihood of incorporating the prior duration into the current reproduction. This sequential bias diminished when the hippocampal activation reached a certain threshold, such as the activation level seen in NR trials – nullifying any significant correlation (Figure 4c). These findings underscore the pivotal role of the hippocampus in mitigating or exacerbating sequential bias.

A logical question arises: why does the hippocampal activation remain elevated in NR trials but not in RR ones? One possibility is that after a trial involving no action, both visual attention and motor readiness are more keenly tuned for the next trial. Indeed, our results revealed higher activity in brain networks associated with executive control and performance monitoring during NR trials compared to RR ones. Recent studies suggest that history biases depend largely on the expectation of making a perceptual decision and the subsequent attention state; when individuals pay more attention to the current stimulus, the influence of the preceding one diminishes (Ceylan and Pascucci 2023; Pascucci et al. 2023). It is worth noting that this sharpened focus on the current stimulus is likely facilitated by the absence of motor activity in the preceding trial. However, we cannot entirely dismiss other factors, such as the frequency distribution of the “Response” and “No-response” trials, as potential contributors. For instance, stimuli from the less frequent “No-response” trials might be more easily disregarded or even cause an opposite effect (Ceylan and Pascucci 2023). Nevertheless, our findings point to more efficient encoding and accurate retention of the current duration during NR trials. This prevents inter-trial memory interference and reduces historical bias. This neural efficiency may reflect a strategic allocation of cognition resources for processing sequential stimuli and optimizing performance (Hsieh et al. 2014; Chanales et al. 2017).

Our findings are broadly consistent with prior studies on non-temporal sequential dependence, corroborating the idea that sequential biases are influenced by the reactivation of the memory trace (Fornaciai and Park 2020b; de Azevedo Neto and Bartels 2021; Ranieri et al. 2022; Sheehan and Serences 2022; Zhang and Luo 2023). For instance, an EEG study employing an auditory pitch categorization task revealed that past information – be it pitch, category choice, or motor response – stores their respective features in memory. These stored features are only reactivated by the corresponding features in the current trial, thereby shifting current neural encoding and giving rise to sequential biases (Zhang and Luo 2023). In the present study, we observed similar dynamics: a negative modulation in the hippocampus by prior reproduction on current encoding, as well as a negative correlation between hippocampal BOLD signals and sequential dependence index. These findings further collectively underscore the crucial role of memory in shaping inter-trial sequential biases.

Turning to future avenues of research, the question of whether hippocampal engagement or associated working memory networks are universally required for sequential dependence in various contexts remains open. Though our study didn’t directly tackle working memory tasks, we did reveal the potential role of working memory in shaping sequential effects in time perception. The relatively long durations (e.g., 1.4 and 1.6 seconds) in our study might place greater demands on memory mechanisms, thereby causing significant differences in neural activations between the NR and RR conditions. Previous literature suggests different neural mechanisms for perceiving sub– and supra-second intervals (Rammsayer 1999; Lewis and Miall 2003; e.g., Hayashi et al. 2014), though supra-seconds are usually longer (e.g., above 3 seconds) than a simple action task that we adopted here. Nevertheless, our results indicated more pronounced differences between NR and RR trials in the hippocampus when longer durations were present in the preceding trial, consistent with increasing evidence highlighting the hippocampus’s central role for longer intervals (Howard and Eichenbaum 2013; Jacobs et al. 2013; Meck et al. 2013; Palombo et al. 2016; Tsao et al. 2022). This is also consistent with the behavioral study on the non-temporal sequential effect (Bliss et al. 2017), which showed that lengthening retention intervals increased the sequential effect.

While it’s clear that the hippocampus plays a key role in sequential-dependent biases in time perception, its role in other types of sequential dependence remains an open question. This complexity mirrors the “sensory recruitment” phenomenon in working memory, where task-specific cortical regions come into play (D’Esposito and Postle 2015). For instance, a recent study used Transcranial Magnetic Stimulation (TMS) to inhibit the left dorsal premotor cortex – a region critical to short-term memory – and observed a significant reduction in serial dependence for judgments of motion speed (de Azevedo Neto and Bartels 2021). On the other hand, when focusing on visual features, evidence from fMRI studies points to the early visual cortex as a key region in sequential dependence (St. John-Saaltink et al. 2016; Sheehan and Serences 2022). Importantly, our findings emphasized that sequential dependence in time perception not only engages specific neural circuits but also relies on the hippocampus’s role in fine-tuning these sequential biases.

In summary, our study revealed that action or not in a preceding trial significantly influences sequential dependence in time perception. When a reproduction task follows passive viewing, both memory and striato-thalamo-cortical networks actively engage, effectively nullifying any sequential biases. In contrast, back-to-back reproduction tasks result in subdued hippocampal activity, which in turn gives rise to prominent sequential biases. Intriguingly, these biases show a negative correlation with individual levels of hippocampal activation. Our findings highlight that sequential biases in time perception do not solely arise at the perceptual stage but also crucially involve the post-perceptual processes. Herr, the hippocampus plays a key role in linking sensory representation to responses.

## Data and code availability statement

The data and analysis code that support the findings of this study will be made available from the author, Si Cheng, upon reasonable request.

## Acknowledgements

This study was supported by German Research Foundation (DFG) research grants SH 166/10-1 to Z.S GL 342/3-2 to S.G. and CH 3093/1-1. The NICUM scanner was financed by the DFG project INST 86/1739-1 (324324095).

## Appendix

### Normalized relative errors

The current reproduction Error (*E_n_*) inherits the general over-/under-estimation bias and the central tendency bias, as well as the sequential bias. Assume the inter-trial sequential bias is independent from the global general and central tendency biases, we can express the following for the reproduction error:

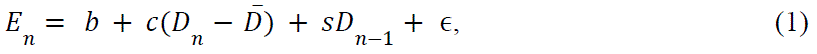

where 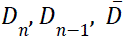 are the current, previous durations, and the mean sample duration, respectively.

The coefficients *b*, *c*, *s* are the general bias, the slope of the central tendency, and the slope of the sequential dependence, respectively. And ɛ is the residual.

When the durations are uniformly distributed and randomly sampled, the conditional distribution of the previous duration on the current duration remains uniform. This means the classical measure of the sequential dependence 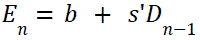 is close to the assumption that the central tendency is linear across the sample durations and averaged out by not considering the current duration. However, this general bias term *b* remains in the equation. To remove the bias term for further analysis, like in fMRI modeling, one approach is to transform the above equation to:

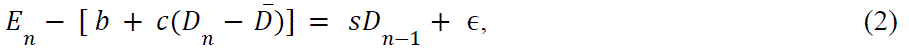

However, this approach has the assumption that the central tendency trend is linear. To relax this assumption, we used another approach, that is, we subtract the mean reproduction from individual sample durations to obtain the relative errors. To equate potential impact of scalar property (i.e.,

Weber scaling), we further normalized the relative error (R*E_n,k_*) as follows:

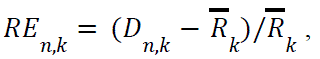

where 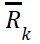 is the mean reproduction of the duration *D_k_*, *D_n,k_* the reproduced duration at trial *n* of a given Duration *D_k_*. When 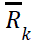 is approximately linear 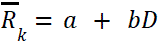, the normalized error is an approximate to Equation (2).

Figure S1 shows the normalized errors were almost flat across all current durations, both for the NR and RR conditions. A repeated-measures ANOVA failed to show any significance (*ps* > 0.084). In contrast, the trends of the normalized error remained similar to the classic measure (Figure 2b), but centered around 0 (see Figure S2). We then calculated the slopes of the linear regression, which was only significantly positive for the RR condition (*b* = 0.054, *t*_(20)_ = 3.011, *p* = .007, *BF_10_* = 6.890), but not for the NR condition (*b* = 0.016, *t*_(20)_ = 1.355, *p* = .190, *BF_10_* = 0.506). Additionally, the slope for the RR condition was significantly larger than that in the NR condition (*t*_(20)_ = 2.213, *p* = .039, *BF_10_* = 1.673, see Figure S3).

**Figure S1.**
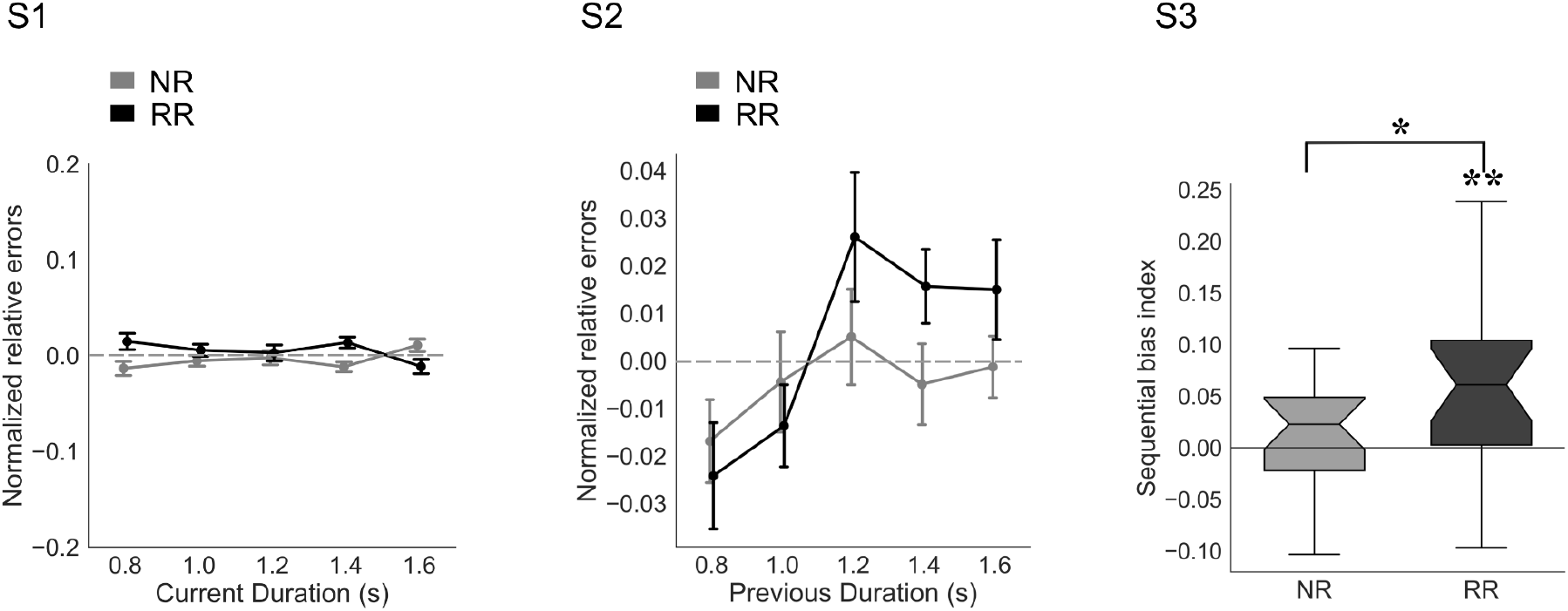
The normalized relative response errors are plotted on the current durations, separated for NR and RR conditions. Error bars represent ± SEM. S2. The normalized relative response errors are plotted on the durations from previous trials, separated for NR and RR conditions. Error bars represent ± SEM. S3. Notched boxplots of the sequential bias for NR and RR conditions. The box plot depicts the slope, measured by the normalized relative errors, for each condition. *p <.05, **p <.01.

## Footnotes

1 During scanning, six participants had large head movements (more than 3 mm of displacement measured by 2nd-degree B-spline interpolation or 3° of rotation in any direction) or distortion in the T1 image. We then replaced with six new participants.

